# The pan-cancer whole-genome landscape of Adolescent and Young Adult solid cancers

**DOI:** 10.64898/2026.10.15.759320

**Authors:** Jeffrey van Putten, Axel Rosendahl Huber, Roelof Koster, Sophie F. Roerink, Joep de Ligt, Paul Roepman, Sjors M. Kas, Ruben van Boxtel, Winette T.A. van der Graaf, Edwin Cuppen, Hilde H. Nienhuis, The GENAYA Consortium

## Abstract

Cancer incidence in adolescents and young adults (AYAs; 18-39 years) is rising, but how their tumors differ genomically from those of older adults (OAs) is unclear. We analyzed paired tumor-normal whole-genomes from 698 AYAs and 5,220 OAs, with subcohorts matched by cancer group to disentangle age from tumor type. AYA tumors carried fewer somatic variants yet largely shared the same genetic drivers; many apparent age differences reflected age-dependent cancer subtypes, such as IDH-mutant glioma and HPV-positive cervical cancer. Colibactin, APOBEC and homologous recombination deficiency-associated signatures were more prevalent in AYA colorectal, gynecological and breast cancers, respectively, but were not more mutagenic when present, suggesting earlier-life activity rather than AYA-specific mutagenesis. Tumor type-concordant pathogenic germline variants were enriched in AYAs (10.9% overall versus 6.6% in matched OAs; p=0.006), often with a somatic second hit. Age thus shaped AYA cancer genomes through mutational processes, subtype composition and germline predisposition, supporting comprehensive tumor-normal profiling beyond driver-focused testing.

## MAIN

Adolescent and young adult (AYA) oncology has emerged as a distinct field in oncology. AYAs, globally mostly defined as diagnosed between 18-39 years, differ from children and older adults in life stage, tumor spectrum, and care needs.^1^ Whereas pediatric cancer care is often concentrated in dedicated centers, AYAs are typically treated across the many adult oncology subspecialties. Cancer incidence is increasing in AYAs.^2^ Some tumor types, such as germ cell tumors and selected sarcomas, peak during the AYA years, whereas others are more characteristic of pediatric or older-adult populations.^2,3^ Nevertheless, systemic treatment strategies for AYAs generally follow tumor-specific pediatric or older adult regimens rather than being informed by age-associated tumor characteristics. Despite recognition of AYAs as a distinct population, it remains unclear how age at diagnosis is reflected in the biology of their cancers.

Age-associated molecular differences have been described for selected tumor types, but the broader genomic landscape of AYA cancers remains partially understood. Existing evidence is fragmented, often tumor-specific, and frequently based on predefined biomarkers or targeted sequencing approaches rather than genome-wide comparisons.^4^ Studies comparing AYAs and other age groups have generally relied on panel-based profiling or small cohorts, while large-scale, pan-cancer comparisons are lacking.^5,6^ Consequently, it remains unclear if specific genomic features distinguish AYA cancers from cancers arising later in life, whether these extend beyond variation in tumor type composition, and which may be relevant for genomics-guided clinical decision-making.

This question is increasingly relevant as molecular biomarkers play a growing role in cancer diagnosis, treatment selection and assessment of cancer predisposition.^7,8^ Current molecular diagnostics primarily assess established tumor-specific biomarkers. They do not, however, enable systematic assessment of broader age-associated genomic patterns. For AYA patients, this leaves uncertainty as to whether molecular findings should be interpreted within pediatric or older adult frameworks, or whether age-associated genomic characteristics provide additional relevant context.

Paired tumor-normal whole-genome sequencing (WGS) enables us to systematically and comprehensively study these differences within a single assay. WGS simultaneously captures somatic small and structural variants, copy-number alterations, gene fusions, viral integrations and mutational signatures, while also enabling assessment of pathogenic germline variants (PGVs).^9,10^ Unlike targeted approaches, which interrogate predefined genomic regions and alteration types, WGS provides a genome-wide, tissue-agnostic framework for studying the full spectrum of genomic variation.^11^ This provides a powerful tool for integrating tumor drivers, mutational processes, and cancer predisposition, and for assessing how these genomic features vary with age.

In this study, we used paired tumor-normal WGS to define the pan-cancer genomic landscape of AYA solid cancers in relation to older-adult cancers. We characterized age-associated differences in somatic mutation burden and driver alterations, placed AYA drivers in the context of pediatric cancers, and assessed PGVs. In addition, we applied refined mutational signature refitting to systematically characterize age-associated mutational processes across tumor types. By integrating these complementary genomic domains, we aimed to identify features relevant to molecular diagnosis, cancer predisposition, and clinical interpretation of tumor genomes in AYA patients.

## RESULTS

The full cohorts consisted of 698 AYAs and 5,220 OAs who underwent WGS as part of routine cancer care or genomic sequencing studies and initiatives. The most frequent cancer groups in the AYA cohort were breast carcinomas (28%), bone and soft tissue tumors (14%), and central nervous system (CNS) neoplasms (12%), whereas the OA cohort consisted mostly of colorectal and intestinal carcinomas (15%), breast carcinomas (13%), and lung carcinomas (13%). Notably, some frequent cancers in OAs such as mesothelioma and kidney carcinomas occurred only sporadically in the AYA cohort, while prostate carcinomas were completely absent in AYAs. Whether patients had received treatments and whether primary or metastatic lesions were biopsied varied by cancer type. (Figure 1)

**Figure 1.**
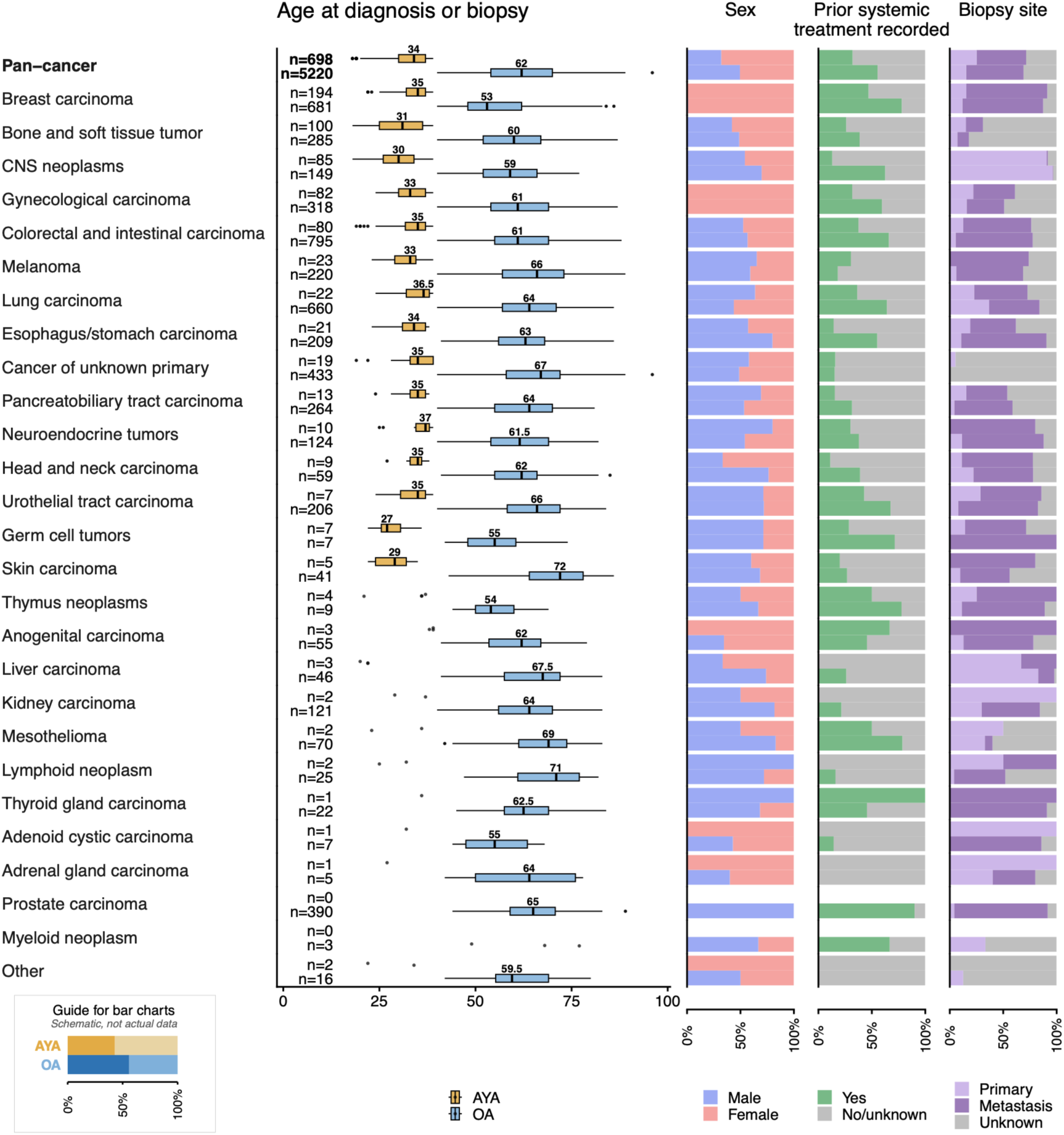
Cohort characteristics of the full study cohort comprising AYAs (n=698) and older adults (n=5,220). Boxplots indicate the interquartile range (IQR), center lines the median, and whiskers the extreme values within 1.5x IQR; median age is indicated as a number above the center line. Outliers are represented by dots. Horizontal stacked bar graphs indicate proportions of patient sex, treatment status before WGS, and whether a biopsy was taken from the primary tumor or a metastatic site.

Because the distribution of major cancer groups differed substantially between AYAs and OAs, pan-cancer genomic comparisons were performed using all 698 AYAs and a 1:1 cancer-group-matched OA cohort (n=698). Cancer-group-specific analyses used all available tumors within each respective cancer group.

### AYA cancers show lower somatic genomic burden but a largely shared driver landscape

We next characterized the somatic genomic landscape of AYA cancers using the cancer-group-matched cohorts. Genome-wide somatic variant burden was consistently lower in AYA tumors across all major variant classes (Figure 2A). AYAs harbored fewer single-nucleotide variants (SNVs; median 7,023 versus 8,676), insertions or deletions (indels; 577 versus 738), multi-nucleotide variants (MNVs; 53.5 versus 68) and structural variant (SV) junctions (201 versus 231; all p<0.001). Homologous recombination deficiency (HRD) was observed in 9.6% of AYA and 7.4% of OA tumors, while microsatellite instability (MSI) was present in 1.4% and 2.9%, respectively.

**Figure 2.**
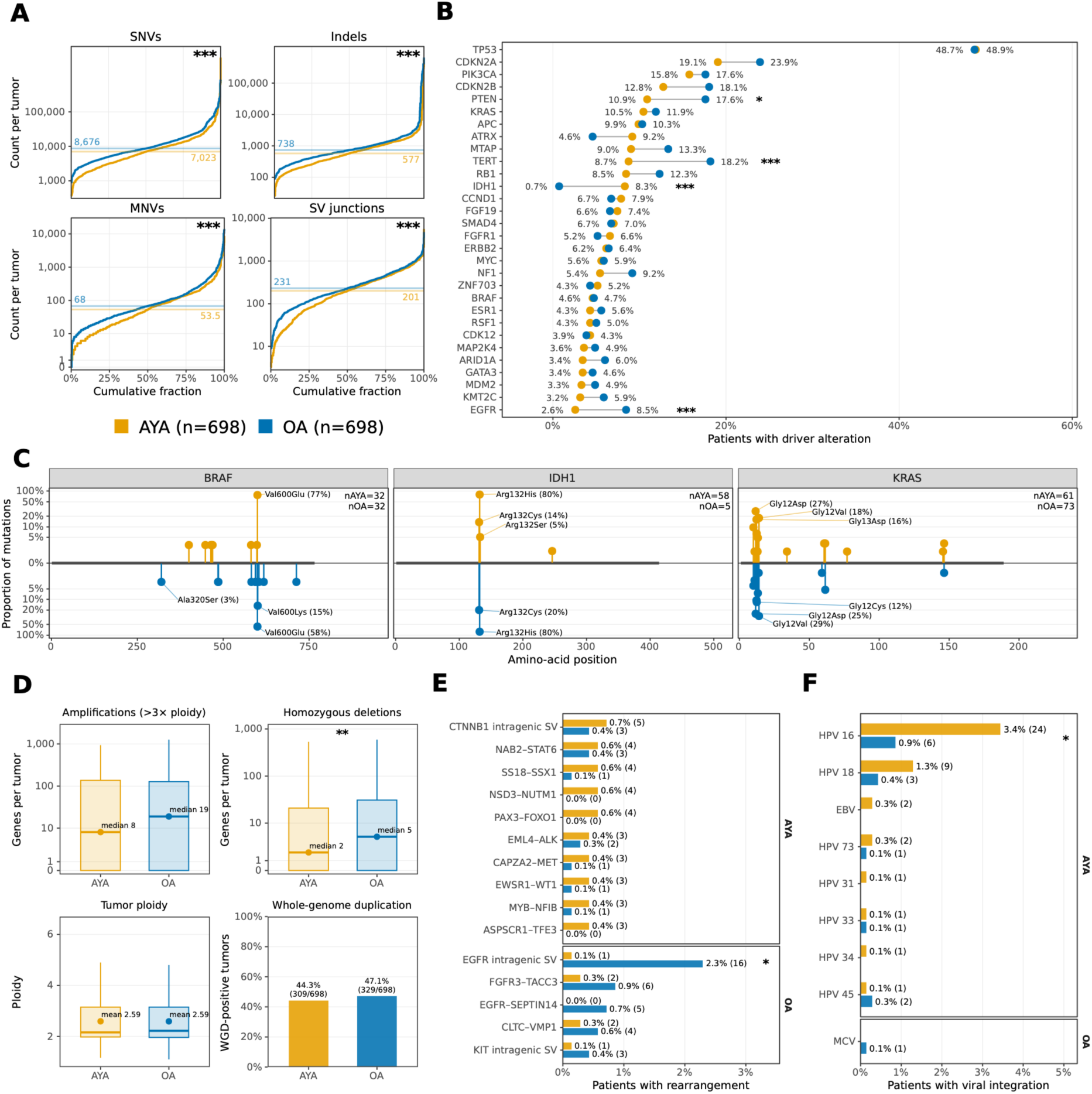
Somatic genomic landscape of AYA cancers in the cancer-group-matched cohort. Pan-cancer comparisons include 698 AYAs and 698 OAs matched by major cancer groups. **A**. Genome-wide burdens of SNVs, indels, MNVs and SV junctions; numbers indicate medians. **B**. Prevalence of recurrent somatic driver alterations comprising mutations, amplifications and deletions, including all genes with a significant age-associated difference and supplemented to a total of 30 genes with the most prevalent remaining drivers across either age group. Gene fusions are shown separately in panel E. **C**. Protein-level variant distribution for selected recurrent drivers, expressed relative to all mutations within each gene and age group; selected recurrent variants are labeled. **D**. Copy-number features, tumor ploidy, and whole-genome duplication. Boxplots indicate the interquartile range (IQR), center lines the median, and whiskers the extreme values within 1.5x IQR. **E**. Prevalence of recurrent gene fusion events, showing the ten most frequent events in AYAs and five additional OA-prevalent events. **F**. Prevalence of viral integrations; integrations detected in AYAs are displayed above OA-only events. Continuous variables were compared using two-sided Wilcoxon rank-sum tests; categorical prevalence differences were assessed using Fisher’s exact tests. Benjamini-Hochberg correction was applied for analyses testing multiple genomic features in B, E and F, whereas p-values in A and D are nominal. * p<0.05, ** p<0.01, *** p<0.001. Abbreviations: SNV, single-nucleotide variant; indel, insertion or deletion; MNV, multi-nucleotide variant; SV, structural variant; HPV, human papillomavirus; EBV, Epstein-Barr virus; MCV, Merkel cell polyomavirus.

To determine whether these differences were also present within cancer groups, we next used all available tumors within each cancer group. Tumor type stratification showed that the lower somatic variant burden in AYAs was broadly maintained, although patterns varied by variant class. Breast cancer was a notable exception: small variant burdens were similar between age groups, but AYA tumors showed a higher SV junction burden (median 438 versus 318; FDR-adjusted p<0.001; Extended Data Fig. 1). This difference remained present within ER-positive disease but was not observed in triple-negative disease. HRD was more frequent in AYA breast cancer (24.9% (48/193) versus 11.1% (75/675); p<0.001). Notably, after stratification by HRD status, SV junction burden remained higher in homologous recombination-proficient AYA breast tumors (median 350 versus 282; p=0.002), indicating that the overall age-group difference was not attributable to the higher prevalence of HRD alone.

**Extended Data Figure 1.**
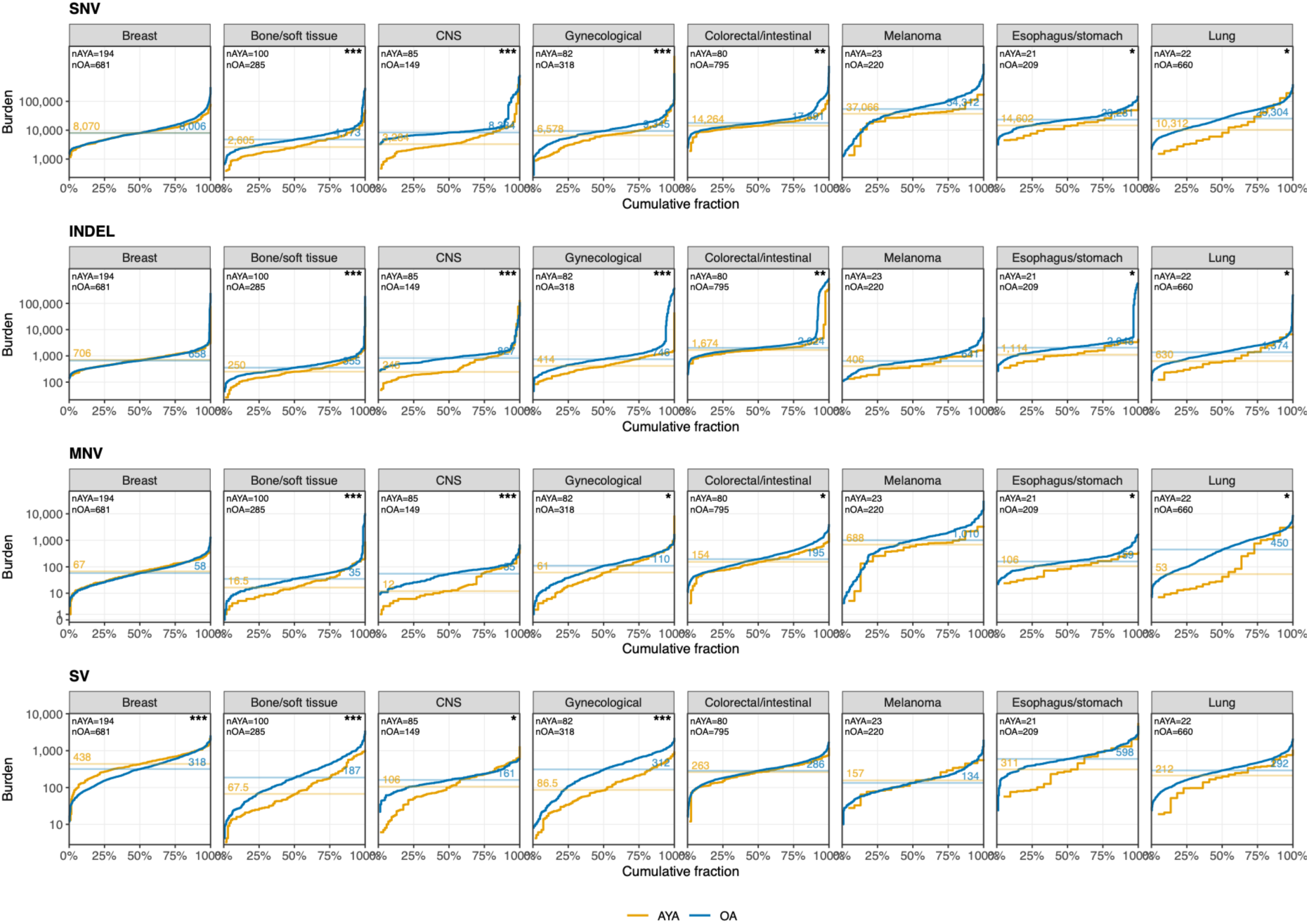
Genome-wide somatic variant burdens stratified by cancer group in the unmatched cohorts. Cumulative distributions are shown for SNVs, indels, MNVs and SV junctions. Numbers indicate median burdens. Asterisks indicate FDR-adjusted significance: * p<0.05, ** p<0.01, *** p<0.001. Abbreviations: SNV, single-nucleotide variant; indel, insertion or deletion; MNV, multi-nucleotide variant; SV, structural variant.

Despite the lower overall somatic genomic burden, the driver landscape was largely shared between groups. *TP53* was the most frequently altered driver gene in both age groups and occurred at nearly identical frequencies, while other recurrent drivers including *CDKN2A*/*B*, *PIK3CA*, *PTEN*, *KRAS* and *APC* were also common to both cohorts. Only selected driver genes retained age-associated differences. In AYAs, *IDH1* alterations were more frequent, whereas *TERT* and *EGFR* alterations occurred more frequently in OAs. (Figure 2B)

Within recurrent driver genes, mutations generally affected the same canonical hotspots, although relative variant frequencies differed for selected genes. For example, *BRAF* alterations in AYAs were almost exclusively Val600Glu (V600E), while OAs showed greater diversity. *IDH1* and *KRAS* event profiles broadly overlapped. (Figure 2C)

Stratification by tumor type showed that recurrent driver profiles were broadly similar between age groups in breast and colorectal cancers, whereas pronounced differences were seen in CNS tumors. (Extended Data Fig. 2A-D, Extended Data Fig. 3A-D) AYA CNS tumors were enriched for *IDH1*, *ATRX* and *TP53* alterations, while *EGFR*, *TERT*, *CDKN2A*/*B*, *MTAP* and *PTEN* alterations predominated in OAs. These patterns were consistent with marked differences in the distribution of molecular glioma subtypes between age groups. (Extended Data Fig. 2C)

**Extended Data Figure 2.**
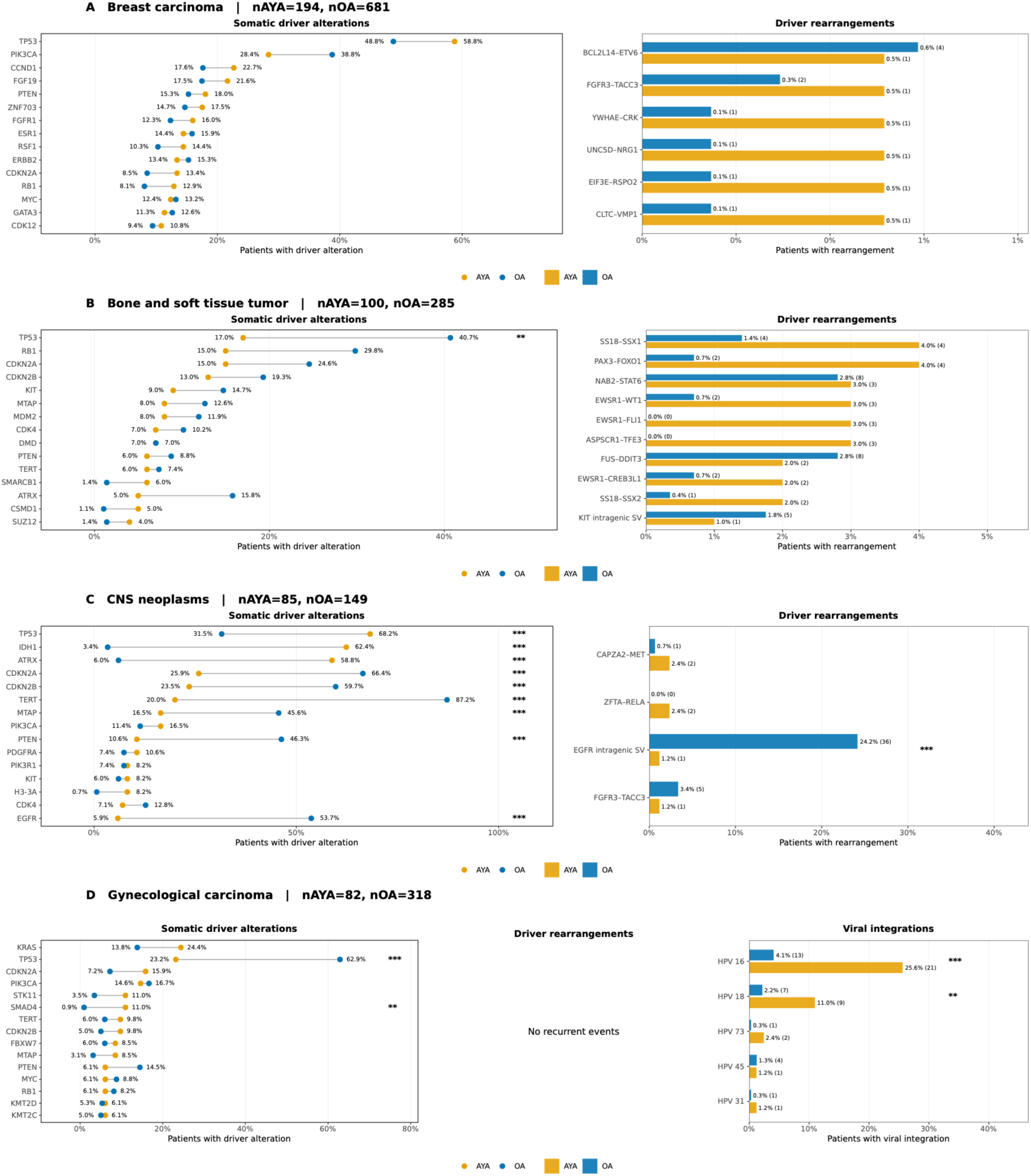
Somatic driver and rearrangement context. Panels display unmatched cohorts and recurrent feature prevalences for (**A**) breast carcinomas, (**B**) bone and soft tissue tumors, (**C**) CNS neoplasms, and (**D**) gynecological carcinomas. Viral integrations are additionally shown for gynecological carcinomas.

**Extended Data Figure 3.**
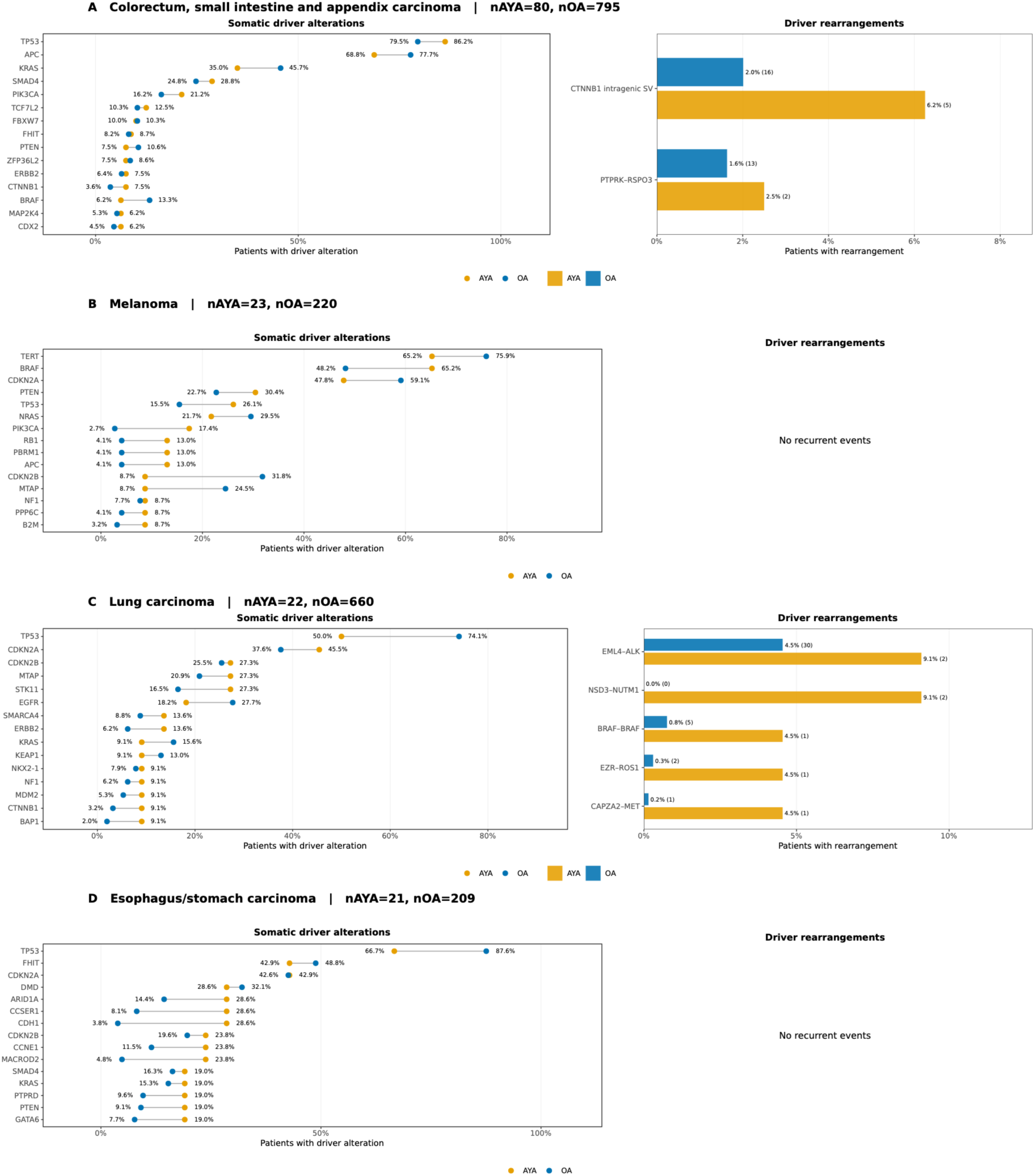
Somatic driver and rearrangement context. Panels display unmatched cohorts and recurrent feature prevalences for (**A**) colorectal, small intestinal and appendiceal carcinomas, (**B**) melanoma, (**C**) lung carcinomas, and (**D**) esophageal and gastric carcinomas.

Beyond individual driver alterations, we assessed broader copy-number architecture. AYA tumors had fewer homozygously deleted genes than matched OAs (median 2 versus 5; p<0.01), whereas amplification burden did not differ significantly. Tumor ploidy was nearly identical (mean 2.59 in both groups), and the prevalence of whole-genome duplication was similar (44.3% versus 47.1%; Figure 2D).

Recurrent gene fusion events also showed a largely shared landscape after cancer-group matching (Figure 2E). No individual recurrent fusion event was significantly enriched in AYAs at the pan-cancer level, whereas intragenic EGFR rearrangements were more frequent in OAs (0.1% versus 2.3%; FDR-adjusted p<0.05). Additional analysis of gene-level disruptive SVs identified no age-associated prevalence differences that remained significant after multiple-testing correction. Within bone and soft tissue tumors, however, combined subtype-defining fusion drivers occurred more frequently in AYAs than in OAs (26.0% (26/100) versus 9.5% (27/285); Fisher’s exact p<0.001; Extended Data Fig. 2B). Within CNS tumors, intragenic *EGFR* rearrangements were markedly more frequent in OAs (24.2% versus 1.2%; FDR-adjusted p<0.001; Extended Data Fig. 2C), reflecting the age-associated differences in molecular subtype composition of CNS tumors.

Viral integrations showed a similar dependence on tumor context (Figure 2F). HPV16 integrations remained more frequent in AYAs after cancer-group matching (3.4% versus 0.9%; FDR-adjusted p<0.05; Extended Data Fig. 2D), whereas other recurrent viral integrations did not differ significantly. The higher HPV16 integration prevalence was concentrated within gynecological carcinoma, where differences in tumor composition remained apparent, since cervical cancers were more frequent among AYAs. Within cervical cancers, HPV integration prevalence did not differ significantly between age groups (97.3% versus 83.9%; Fisher’s exact p=0.085).

The corresponding pan-cancer genomic landscape using all 5,220 unmatched OA tumors is shown in Extended Data Fig. 4. Compared with the cancer-group-matched analysis, the full cohort showed broader differences across driver alterations, copy-number features, rearrangements and viral integrations. Several of these differences were attenuated after matching, whereas differences in genome-wide somatic variant burden and selected genomic features persisted. This illustrates the substantial contribution of cancer-group composition to pan-cancer age-associated genomic differences.

**Extended Data Figure 4.**
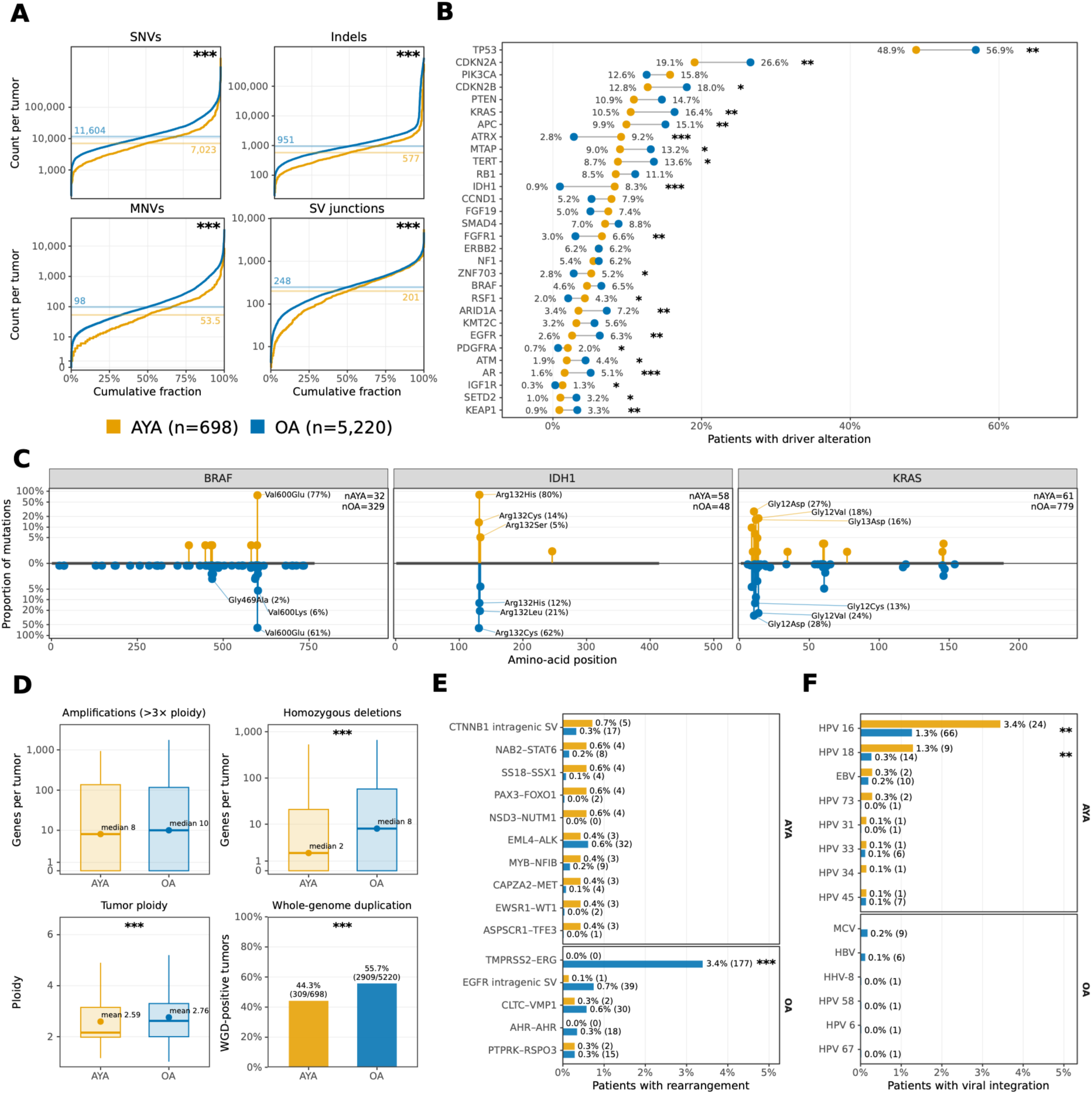
The somatic genomic landscape of AYA cancers compared with all 5,220 OA cancers. Cohorts were not matched by cancer group.

Together, these findings indicate that cancer in AYAs is characterized by a lower overall burden of somatic genomic alterations, but largely shares the established recurrent oncogenic driver landscape observed in OAs. Many of the most pronounced age-related differences in individual genomic features reflected differences in tumor type and molecular subtype composition. Overall, age-associated differences were more pronounced in genome-wide mutational burden and molecular context than in recurrent oncogenic drivers.

### Comparison with pediatric cancer drivers

To place the AYA driver landscape in a broader age context, we compared gene-level driver prevalence with pediatric cancers that are also prevalent in AYAs. We derived driver prevalences of soft tissue and CNS tumors from the St. Jude PeCan resource.

In soft tissue tumors, distinct age-associated driver patterns were observed.

*SMARCB1* and *PAX3* were the most prevalent in pediatric tumors (18% and 13%, respectively) and became progressively less frequent in AYA and OA tumors. Among AYAs, several partner genes involved in fusion-driven sarcoma subtypes were among the most prevalent, including *EWSR1* (10%), *SS18* (7%), and *TFE3* (6%). Notably, *NF1* and *CREBBP* showed lower prevalence in AYAs than in both pediatric and OA tumors. In OAs, *TP53, RB1, CDKN2A/B* and *ATRX* were among the most prevalent drivers. (Figure 3A)

**Figure 3.**
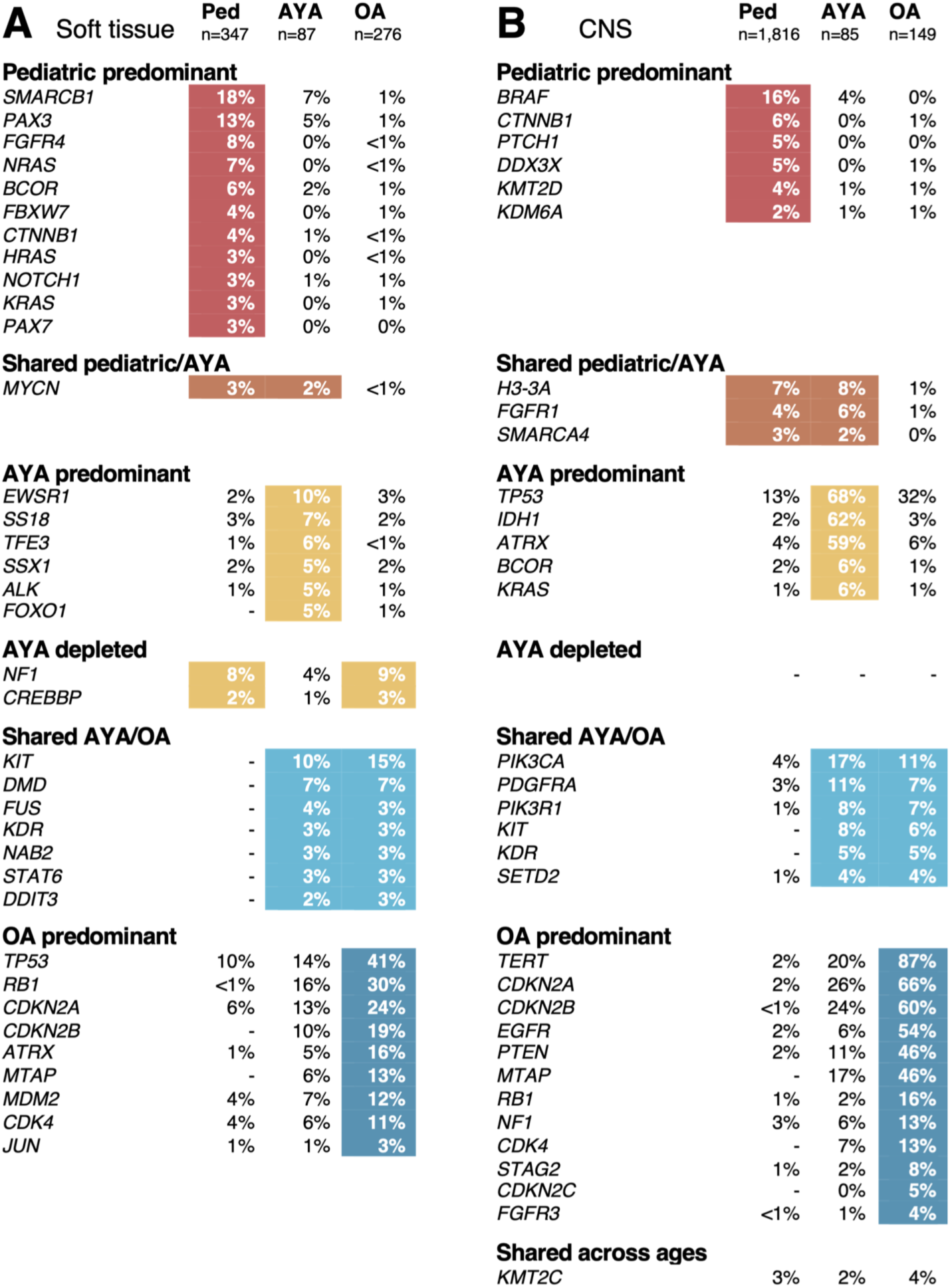
Comparative prevalence of recurrent driver gene alterations in pediatric, AYA and OA soft tissue and CNS tumors. Recurrent driver gene alterations for (**A**) soft tissue tumors and (**B**) CNS tumors. Genes represent the union of the 20 most prevalent driver genes in each age group and are grouped according to their age-associated prevalence patterns, illustrating pediatric-predominant, shared pediatric/AYA, AYA-predominant, AYA-depleted, shared AYA/OA, OA-predominant, and shared across ages. Age groups were classified as sharing a predominant pattern when their prevalence was within 1.5-fold of the prevalence in an adjacent age group. Pattern assignment used unrounded prevalence estimates for AYAs and OAs and whole-number prevalences reported by the St. Jude PeCan resource for pediatric tumors; percentages shown in the figure are rounded to whole numbers. AYA and OA gene-level prevalence were calculated from somatic driver mutations, amplifications, deletions and gene fusions. Comparisons are descriptive because the PeCan and Hartwig datasets use different analytical frameworks. A dash indicates that the gene was not reported as a driver in the corresponding dataset.

In CNS tumors, the driver landscape showed distinct age-associated patterns. *BRAF* alterations were the most frequent in pediatric brain tumors (16%), where they consisted mainly of fusions, while this was markedly lower among AYAs (5%) and absent in OAs. *H3-3A and FGFR1* alterations occurred at comparable frequencies in pediatric and AYA brain tumors, but were rare in OAs. Conversely, *IDH1*, *ATRX*, and *TP53* were predominant in the AYA cohort compared with both pediatric and OA tumors. Other alterations, including *TERT* and *CDKN2A*/*B*, increased across pediatric, AYA and OA tumors. (Figure 3B)

Together these comparisons show that the AYA driver landscape comprises features shared with both pediatric and OA cancers but also includes features that were particularly prevalent during the AYA age, and reflects age-associated differences in molecular subtype composition.

### Mutational processes underlying tumor etiology

We refitted COSMIC single-base substitution (SBS), doublet-base substitution (DBS) and indel (ID) signatures to the mutation profiles of our cohort to obtain insights into the biological processes shaping AYA and OA cancers.^12^ Reconstructed signature profiles using *SigProfilerAssignment* showed high cosine similarity with the observed mutational spectra in both age groups, indicating that COSMIC reference signatures adequately captured the mutational profiles of AYA and OA cancers.^12^ (Extended Data Fig. 5).

**Extended Data Figure 5.**
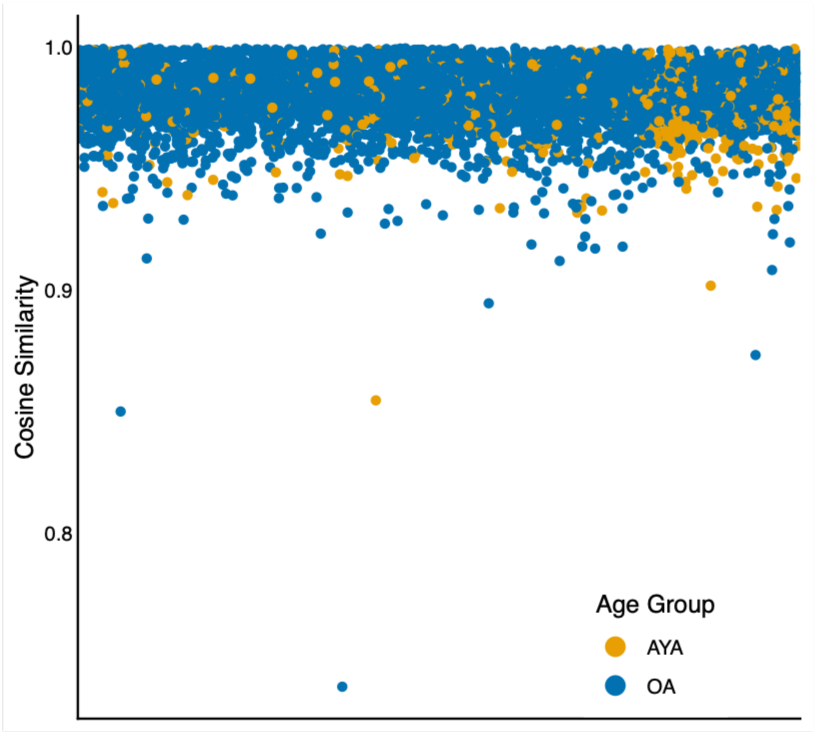
Cosine similarity of all patients in the cohort following refitting of COSMIC signatures. High cosine similarity indicates matching of the signature contributions with observed mutational patterns. One sample with a cosine similarity to the reconstructed profile of <0.8 was observed, which was previously described and identified as a psoralen-induced basal cell carcinoma (Olafsson et al, Nat Genet, 2023).

Upper panels in D, E and G show signature-attributed mutation counts among signature-positive tumors; lower panels show signature prevalence. Abbreviations: SBS, single-base substitution; ID, insertion or deletion; DBS, doublet-base substitution; NER, nucleotide excision repair; DSB, double-strand break; HPV, human papillomavirus.

### Age-associated differences reflect both prevalence and signature contribution

Age-associated signature differences may reflect a higher prevalence or longer exposure of mutational processes, a greater mutational burden when present, or both. Consistent with cumulative age-related mutagenesis, clock-like signatures SBS1, SBS5, and SBS40a showed lower contributions in AYAs than in OAs across several cancer types. (Figure 4A-C)

**Figure 4.**
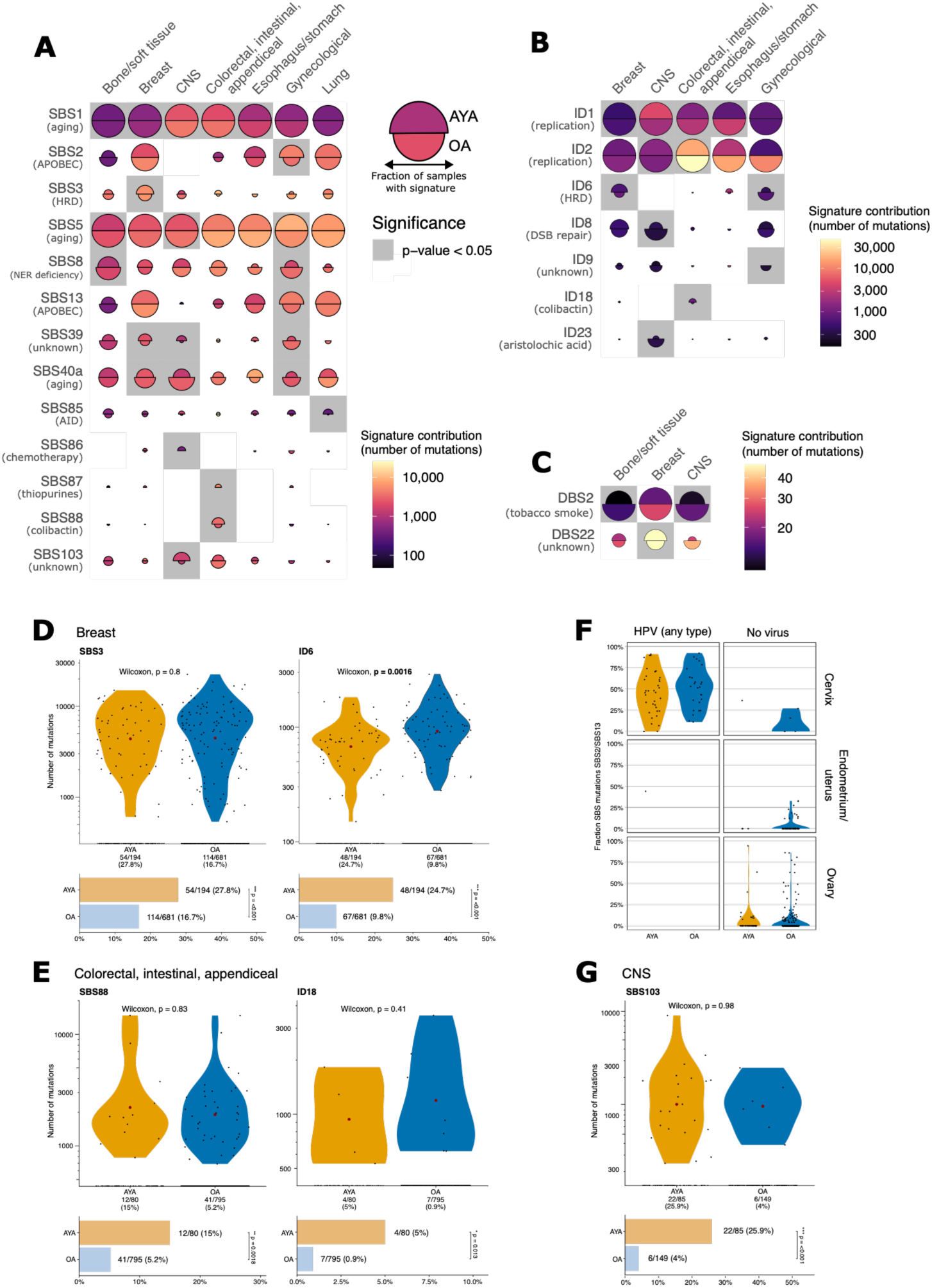
Mutational signature landscapes and selected age-associated processes. **A-C.** SBS, ID and DBS signature contributions in AYA and older-adult tumors across cancer groups after *SigProfilerAssignment* refitting. Split orbs display fractions of AYAs and OAs within cancer type groups, with AYAs shown above and OAs below in each orb panel. Gray backgrounds indicate significant age-group differences (FDR-adjusted p<0.05). Proposed etiology of signatures is based on COSMIC. **D**. Prevalence and contribution of SBS3 and ID6 in breast carcinomas. **E**. Prevalence and contribution of SBS88 and ID18 in colorectal, small intestinal and appendiceal carcinomas. **F**. Fraction of SBS mutations attributed to SBS2/SBS13 in gynecological carcinomas according to tumor site, age group and HPV integration status. **G**. Prevalence and contribution of SBS103 in CNS neoplasms.

HRD-associated mutagenesis was more prevalent in AYA breast cancer, reflected by a higher prevalence of SBS3 and ID6. SBS3 was present in 27.8% of AYA tumors compared with 16.7% of OAs (Fisher’s exact p<0.001), whereas the number of mutations attributed to SBS3 did not differ among SBS3-positive tumors (Wilcoxon p=0.80). Similarly, ID6 was more prevalent in AYAs (24.7% versus 9.8%, Fisher’s exact p<0.001), although among ID6-positive tumors its contribution was higher in OAs (Wilcoxon p=0.0016). These findings indicate that the enrichment of HRD-associated mutagenesis primarily reflected a higher prevalence of these processes, although for ID6 this was accompanied by a lower mutational contribution in AYAs when present. (Figure 4D)

A similar prevalence pattern was observed in CRC. Colibactin-associated mutagenesis was more prevalent in AYA CRC, consistent with previous observations.^13,14^ SBS88 was detected in 15.0% of AYAs compared to 5.2% of OAs (Fisher’s exact p=0.0018), and ID18 in 5.0% versus 0.9%, respectively (Fisher’s exact p=0.013). Among signature-positive tumors, however, the number of mutations attributed to either signature did not differ between age groups (SBS88, Wilcoxon p=0.83; ID18, Wilcoxon p=0.41; Figure 4E). This pattern is compatible with colibactin-induced mutagenesis being an episodic rather than a continuously active process, with a greater proportion of AYA CRC showing evidence of exposure, but a comparable mutational contribution among signature-positive tumors.

### Tumor and molecular context explain selected AYA-enriched signatures

An additional exposure-related signal was observed in AYA CRC. SBS87, a signature previously associated with thiopurine exposure, was detected in three AYAs and no OAs with CRC. (Figure 4A) Two of these patients had documented inflammatory bowel disease and prior immunosuppressive treatment, although previous thiopurine exposure could not be confirmed from available treatment records. Non-oncological medical history was unavailable for the third patient, limiting further interpretation of this association.

APOBEC-associated mutagenesis appeared enriched in AYA gynecological cancers, reflected by higher contributions of SBS2 and SBS13. Downstream analyses showed that high SBS2/SBS13 contributions were concentrated predominantly in human papillomavirus (HPV)-positive cervical cancers (Figure 4F). Among cancers with HPV integration, APOBEC-associated mutational activity was not higher in AYAs and was instead greater in OAs. Most HPV integrations involved HPV16 or HPV18 and no associations were seen in HPV-positive head and neck or anogenital cancers. The enrichment of SBS2 and SBS13 across AYA gynecological cancers therefore appears to reflect the greater representation of cervical cancers as well as HPV integration in the younger cohort, rather than a general age-related increase in APOBEC mutagenesis.

A similar compositional effect was observed for SBS103, a signature with currently unknown etiology.^14^ SBS103 was detected in 25.9% of AYA CNS tumors compared with 4.0% of OA CNS tumors (Fisher’s exact p<0.001), whereas its contribution among SBS103-positive tumors did not differ between age groups (Wilcoxon p=0.98; Figure 4G). Further molecular classification showed that SBS103 was strongly concentrated in *IDH*-mutant glioma, which is consistent with previous observations in CNS tumors,^15^ particularly in tumors molecularly classified as *TP53*-mutated astrocytoma. OA astrocytomas also harbored SBS103, indicating that this signature is not AYA-specific and that its apparent enrichment in AYAs largely reflects differences in CNS molecular subtype composition occurring at different ages.

Together, these findings show that age-associated mutational signature differences reflect a combination of cumulative age-related mutagenesis, differences in prevalence of specific mutational processes, and differences in tumor and molecular subtype composition between younger and older patients.

### Germline cancer predisposition is enriched in AYAs

Among 698 AYAs, 105 (15.0%) carried at least one PGV in a cancer predisposition gene. Because the clinical relevance of germline predisposition is strongly dependent on tumor type, we next classified PGVs according to their concordance with the tumor type in which they were found and compared AYAs with cancer group-matched OAs.

PGVs associated with the tumor type in the respective patient were more frequent in AYAs than in OAs (10.9% versus 6.6%, Fisher’s exact p=0.006), whereas the prevalence of PGVs in genes not associated with the tumor type was similar between age groups (4.2% versus 5.6%, Fisher’s exact p=0.26). The higher prevalence of germline predisposition in AYAs was therefore concentrated in variants with an established relationship to the cancer in which they were identified (Figure 5A).

**Figure 5.**
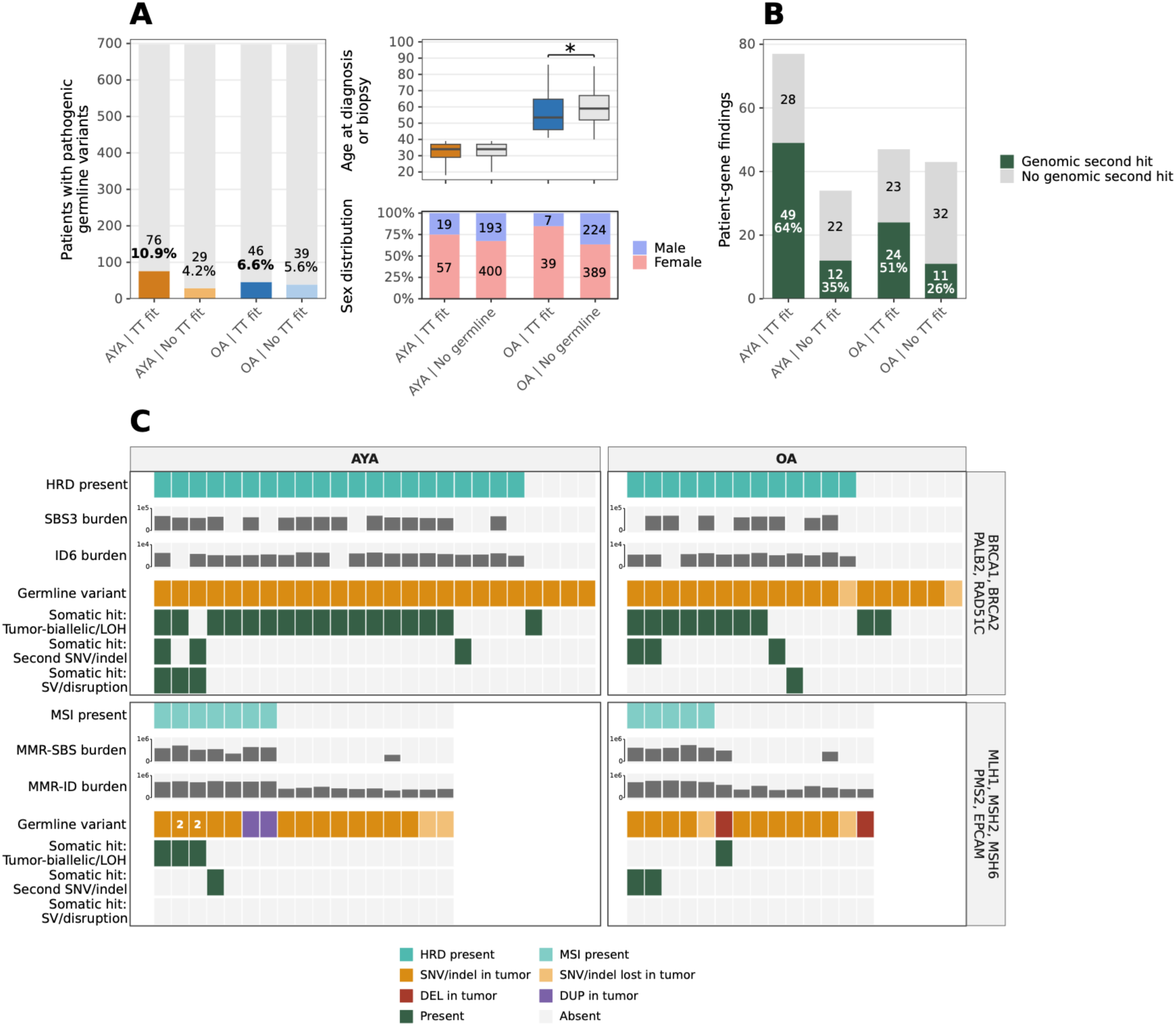
Germline events and tumor genomic context. **A**. Prevalence of PGVs in 698 AYAs and 698 cancer group-matched OAs, stratified according to whether the affected gene was associated with the patient’s tumor type (tumor type-concordant; TT fit) or not (No TT fit) based on the guideline used in the Netherlands.^16^ Numbers and percentages indicate patients in each category. Age distribution and sex distribution are shown for patients with tumor type-concordant germline variants and patients without a PGV. The asterisk denotes p<0.05 by Wilcoxon rank-sum test. **B**. Presence of a genomic second hit among tumor type-concordant and non-concordant pathogenic germline findings in AYAs and OAs. Numbers indicate patient-gene findings, with each gene counted once per patient; percentages indicate the proportion accompanied by a genomic second hit. **C**. Integrated germline, somatic and tumor genomic profiles for carriers of PGVs in homologous recombination repair (HRR; *BRCA1*, *BRCA2*, *PALB2* and *RAD51C*) or mismatch repair (MMR; *MLH1*, *MSH2*, *MSH6*, *PMS2* and *EPCAM*) genes, shown separately for AYAs and OAs. Columns represent individual patients. For HRR germline carriers, rows indicate CHORD-defined HRD status, SBS3 and ID6 burden, the germline alteration and genomic second-hit mechanisms. For MMR germline carriers, rows indicate *MSIsensor*-based MSI status, MMR-associated SBS and ID burden, the germline alteration and genomic second-hit mechanisms. The germline track indicates whether the pathogenic germline SNV or indel was detected in the tumor. A number 2 within the germline track indicates two PGVs within the same gene. Genomic second hits comprise tumor-biallelic loss or LOH, a second SNV or indel, or a structural variant affecting the same gene; because allelic phasing was generally unavailable, these were considered putative. The tumor-biallelic/LOH track also displays biallelic germline *PMS2* alterations, corresponding to two CMMRD cases. Abbreviations: DEL, deletion; DUP, duplication; LOH, loss-of-heterozygosity.

Among the most frequently affected cancer predisposition genes in both age groups were genes involved in homologous recombination repair (HRR) and mismatch repair (MMR), while alterations in genes such as *SDHA*, *APC* and *TP53* were particularly common in AYAs (Extended Data Fig. 6). Among OAs, tumor type-concordant germline variants were associated with younger age and female sex, whereas no age or sex association was observed among AYAs (Figure 5A).

**Extended Data Figure 6.**
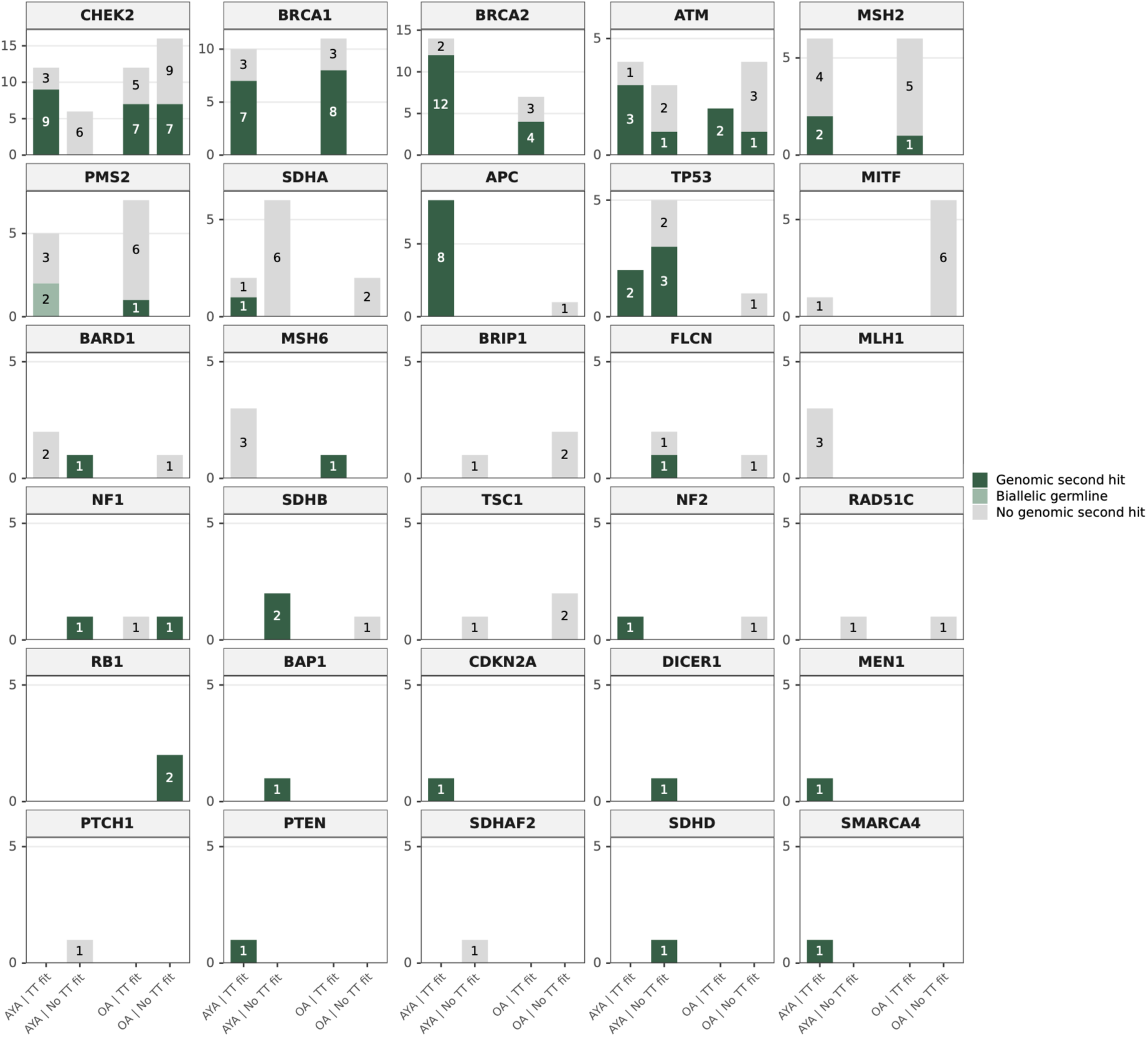
Gene-level distribution of pathogenic germline variants and genomic second hits. PGVs are shown by affected cancer predisposition gene and stratified by age group and concordance with the patient’s tumor type (tumor type-concordant, TT fit; non-concordant, No TT fit). Stacked bars indicate patient-gene findings with or without a detectable genomic second hit, or biallelic germline alterations, with each gene counted once per patient and numbers indicating finding counts. Genes are ordered by overall frequency of pathogenic germline findings. A genomic second hit was defined as tumor-biallelic loss or LOH, an additional somatic SNV or indel, or structural variant of the affected gene.

We next evaluated whether PGVs were accompanied by somatic events, referred to here as genomic second hits. These comprised loss-of-heterozygosity (LOH) or tumor-biallelic loss, additional SNVs or indels, and SVs affecting the same gene; as allelic phasing relative to the germline variant was generally unavailable, these were considered putative.

Among tumor type-concordant findings, a genomic second hit was identified in 63.6% of AYA and 51.1% of OA findings, with no significant difference between age groups (Fisher’s exact p=0.19). Within both age groups, however, genomic second hits were more frequent among tumor type-concordant than non-concordant findings (AYA: 63.6% versus 35.3%, Fisher’s exact p=0.007; OA: 51.1% versus 25.6%, p=0.017), supporting the biological relevance of tumor type-concordance (Figure 5B).

At the patient level, 7.0% (49/698) of AYAs compared with 3.3% (23/698) of OAs carried a tumor type-concordant PGV accompanied by at least one detectable genomic second hit (Fisher’s exact p=0.002). One of these 23 OAs carried tumor type-concordant germline variants in two different genes, both with a genomic second hit. Although the probability of detecting a genomic second hit among tumor type-concordant germline findings was comparable between age groups, the combination of relevant germline predisposition and additional tumor genomic features was more frequent in AYAs. Gene-level analyses showed variation in genomic second hit prevalence across cancer predisposition genes (Extended Data Fig. 6).

### Somatic genomic context for HRR and MMR germline alterations

Whole-genome data additionally enabled independent assessment of the downstream functional consequences of cancer predisposition through characteristic genome-wide patterns of DNA repair deficiency. We therefore integrated germline and somatic alterations in recurrent HRR and MMR genes with WGS-based classifications of HRD and MSI.

Among carriers of PGVs in HRR genes, genomic second hits were identified in 76% (19/25) of AYAs and 63% (12/19) of OAs. Independently, CHORD classified 84% (21/25) of AYAs and 68% (13/19) of OAs as HRD-positive. HRD-associated SBS3 and ID6 contributions were concordant with CHORD classification and provided additional evidence of the associated mutational scarring in HRD.

Among MMR germline carriers, genomic second hits were less frequently detected (AYA: 12% (2/17); OA: 21% (3/14)), whereas MSI was identified independently by the pipeline’s MSIsensor-based classifier in 41% (7/17) and 36% (5/14), respectively. Notably, two AYAs with CNS tumors carried compound heterozygous pathogenic germline *PMS2* variants, consistent with constitutional mismatch repair deficiency (CMMRD). MMR-associated SBS and ID burdens were higher in MSI tumors in both age groups, although lower-level contributions, particularly of ID1 and ID2, were also observed in MMR carriers with MSS phenotype. These integrated profiles showed similar patterns across age groups: HRR germline variants were frequently accompanied by both genomic second hits and CHORD-defined HRD, whereas MSI was often observed in MMR germline carriers without a detectable genomic second hit. In some cases, including the CMMRD cases, pathway inactivation could be explained by biallelic germline predisposition. In other cases, inactivation may be caused by alternative mechanisms not captured by the genomic second hit analysis, such as epigenetic silencing. (Figure 5C)

Together, these findings indicate that tumor type-concordant cancer predisposition is more frequent in AYAs, while the downstream tumor biology associated with such predisposition, including genomic second hits and DNA repair deficiency, is broadly similar across age groups.

## DISCUSSION

This study provides a comprehensive whole-genome characterization of cancers arising in adolescents and young adults, an age group positioned between pediatric and older adult oncology. Although AYAs are recognized as a distinct clinical population, particularly because of their age-specific care needs, it has remained unclear which genomic characteristics distinguish cancers arising in AYAs from those in older adults, and to what extent these reflect tumor and molecular subtype composition. Our findings do not support a uniform AYA-specific molecular architecture. Rather, age at diagnosis is reflected across multiple genomic layers, including in somatic mutation burden, the distribution of molecular cancer subtypes, mutational processes, and cancer predisposition.

Previous studies have described age-associated genomic differences using panel-based sequencing or WGS in small cohorts.^5,6^ Our pan-cancer analysis extends these observations to a much larger cohort of patients whose tumors all underwent WGS. Pan-cancer comparisons after matching showed that several age-associated differences were attenuated, whereas others were maintained. This distinction highlights two complementary aspects of the AYA genomic landscape: differences arising from the distinct spectrum of cancers occurring at younger ages, and genomic differences that remain after accounting for major cancer-group composition.

Despite a largely shared recurrent driver landscape, several signals only became visible when broader genomic features or molecular context were considered. One example is somatic mutational burden, which was generally lower in AYAs across cancer types, while breast cancer formed a notable exception for SVs. The higher SV junction burden in AYA breast cancer co-occurred with greater prevalence of HRD, consistent with previous studies,^17,18^ although its persistence among homologous recombination-proficient tumors indicates that HRD does not fully account for this difference. The molecular heterogeneity observed in CNS tumors and fusion-driven sarcomas, which have different occurrence patterns across age groups,^19–21^ further illustrates how broad tumor categories can comprise molecular subtypes with strongly age-dependent distributions. Comparison with pediatric cancer drivers showed that AYAs are not simply intermediate between childhood and older adulthood: some molecular features were shared with adjacent age groups, whereas others were most prevalent or even depleted during the AYA years.

Whole-genome analysis also allowed us to study mutational processes beyond recurrent cancer drivers. Clock-like signatures showed lower contributions in AYAs, consistent with cumulative mutagenesis with increasing age. In contrast, several processes enriched in AYAs reflected a greater proportion of signature-positive tumors rather than higher signature-attributed mutation counts. Colibactin-associated mutagenesis in CRC, for instance, was notably more prevalent in AYAs but displayed similar levels of signature-attributable mutations when present. This observation supports and expands previous observations in early-onset CRC.^13,14^ The apparent enrichment of APOBEC-associated mutagenesis across AYA gynecological cancers further demonstrated how signature patterns that appear age-associated may instead arise from differences in tumor site, molecular subtype composition, or, in this case, HPV-related tumor context. SBS103, whose underlying etiology remains unknown, was enriched in AYAs and concentrated in *TP53*-mutated astrocytoma. However, it was also observed in OA tumors and has previously been reported in CRC populations,^14^ further illustrating this complexity. Its apparent association with younger age may reflect the age-dependent distribution of molecular subtypes, although whether age also influences the contribution of mutational processes underlying these subtypes requires further study. These findings underline the importance of interpreting mutational signatures within their tumor context rather than as isolated age-specific processes. Age-associated differences in mutational signatures of unknown etiology observed here warrant future correlation with clinical and environmental exposures, providing opportunities to study potential etiologic factors associated with AYA cancer development.

Cancer predisposition presented another genomic layer associated with age. An increased contribution of cancer predisposition has previously been reported in younger patients across various tumor types.^6,22–25^ Here, we found that AYAs harbored significantly more PGVs in genes with an established association with the respective tumor type than OAs (10.9% versus 6.6%), whereas PGVs without such tumor type-association occurred at similar frequencies in both age groups. This suggests enrichment of clinically plausible cancer predisposition in AYAs. Within the OA cohort, tumor type-concordant PGVs were concentrated at younger ages. For context, previous non-age-stratified pan-cancer whole-genome analyses of metastatic solid tumors identified PGVs in 9.0% and 7.9% of all adults.^10,26^ At patient level, AYAs more often carried a detectable genomic second hit accompanying a tumor type-concordant PGV. Integration of germline alterations with independently derived HRD and MSI classifications further showed that downstream tumor biology associated with inherited DNA damage repair defects was broadly similar across age groups. However, because the OA comparison cohort was matched to the AYA tumor spectrum, these findings are related to age-associated predisposition within the tumor types represented among AYAs and may not represent the overall contribution of germline predisposition to all cancers in OAs.

Taken together, AYA cancers largely share the recurrent driver landscape of OAs, while age-associated differences persist in mutational burdens, molecular subtype composition, mutational processes and cancer predisposition. Several AYA-enriched mutational processes were also observed in OAs and differed primarily in prevalence rather than the extent of mutagenesis, consistent with contributions earlier in life rather than AYA-specific mechanisms.

Our cohort only included patients whose tumors underwent WGS, and several cancers common in the broader AYA population, including testicular tumors, thyroid cancers and MSI colorectal cancers, were underrepresented, limiting generalizability. Nevertheless, these findings may have implications for genomic diagnostics in AYAs. Tumor type remains the primary framework for interpreting recurrent drivers, more so than age. However, broader genomic profiling can reveal additional age-associated context that conventional driver-focused profiling may miss. In pediatric oncology, whole-genome and transcriptome sequencing are increasingly embedded into routine diagnostics,^27,28^ while paired tumor-normal WGS has also demonstrated clinical utility in adult oncology more broadly.^29^ By jointly capturing somatic drivers, structural variants, mutational processes and inherited predisposition, WGS may be particularly informative for AYAs beyond recurrent driver assessment, while also providing a genomic backbone for multi-omic precision oncology.

## ACKNOWLEDGEMENTS

We thank all patients whose genomic and clinical data contributed to this study. We thank the staff of Hartwig Medical Foundation involved in WGS diagnostics and the data infrastructure supporting the generation and curation of these data. We further thank Francisco Martínez-Jiménez, Joseph Usset, Natasa Mortvanski, David Koetsier, Erik van Dijk, Yannick Bijl, Hans Clevers and the Clevers lab for methodological and computational input. Patient and tumor characteristics as well as survival data were obtained from the Netherlands Cancer Registry (NKR) which is maintained by the Netherlands Comprehensive Cancer Organization (IKNL).

## AUTHOR CONTRIBUTION STATEMENT

JvP had access to all data for the present study and all analysis outputs and takes responsibility for the integrity of the data and the accuracy of the overall analysis. JdL and SFR enabled linkage to external clinical data sources; RK performed clinical interpretation and expert review of the germline data before analysis. JvP, EC and HN developed the methodology; JvP performed the formal analyses and created the data visualizations for the somatic landscape and germline analyses; ARH carried out the methodology, performed the formal analyses, and created the data visualization of the mutational signature refitting and subsequent analyses. JvP wrote the original manuscript under supervision of HN and EC; ARH drafted the methods describing the mutational signature analyses and contributed to the corresponding results and interpretation; RK contributed to the germline-related sections. JdL, SFR, PR, SK, RvB and WvdG provided critical feedback on the manuscript. All authors reviewed and approved the final manuscript.

## The GENAYA Consortium

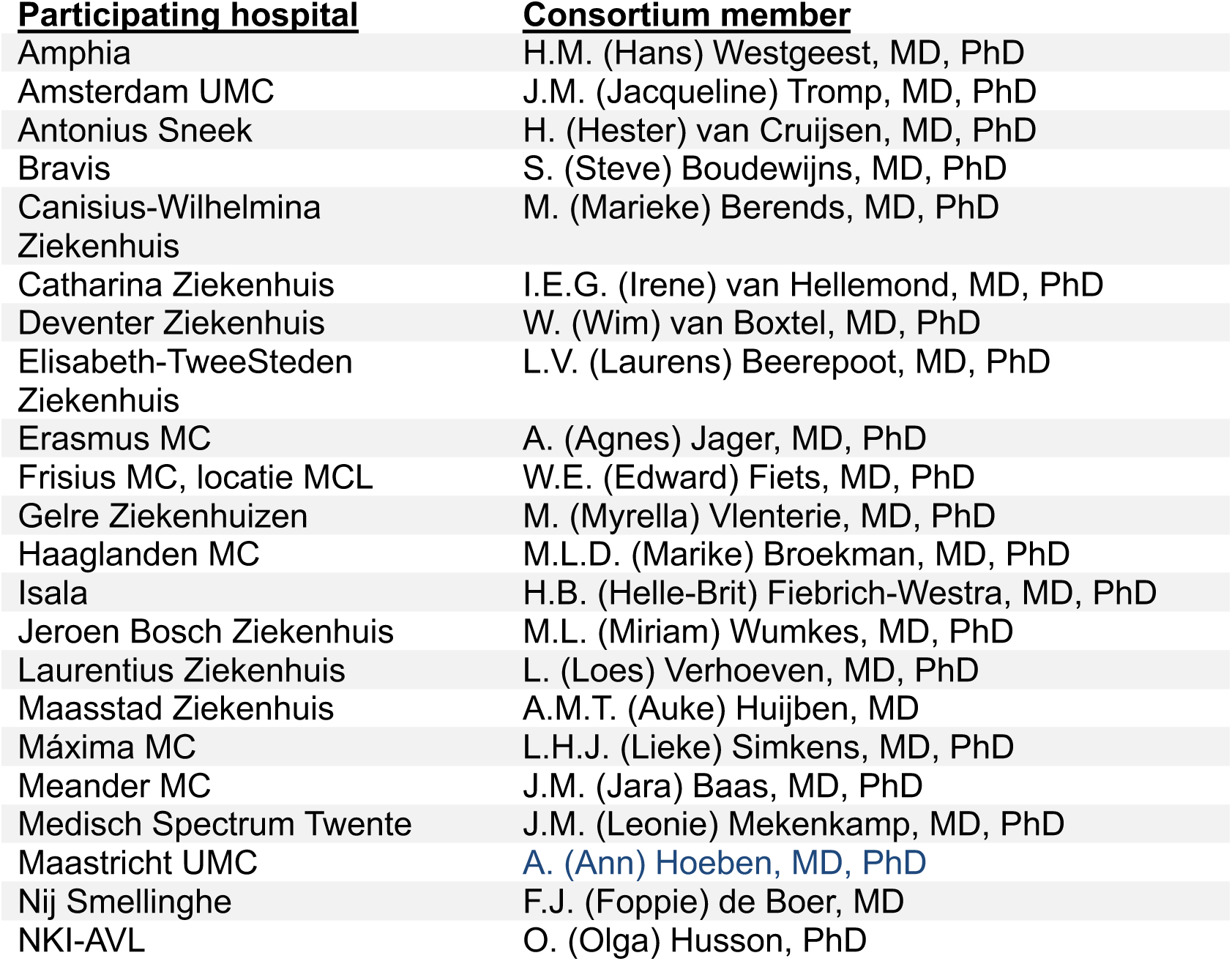
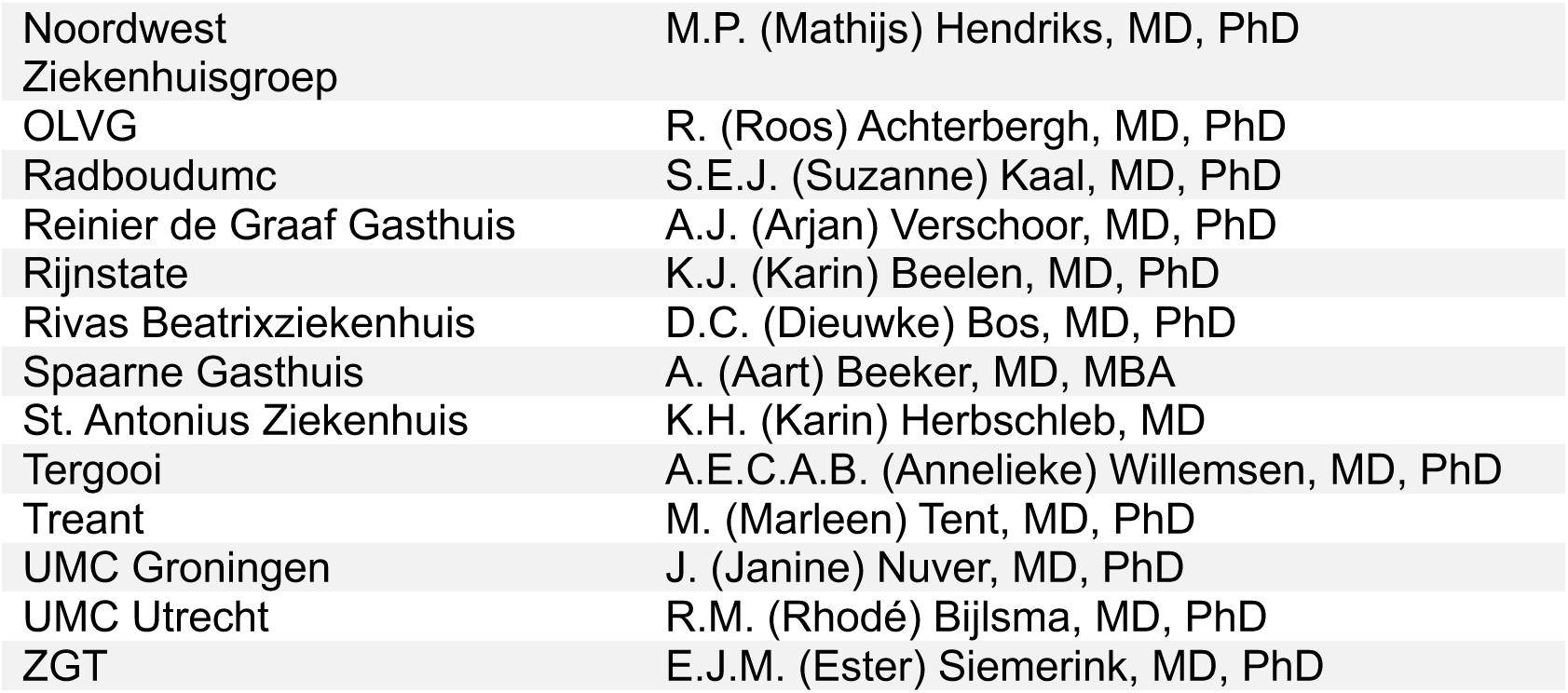

## METHODS

### Study design and cohort definition

This retrospective pan-cancer study included patients with solid cancers who underwent paired tumor-normal whole-genome sequencing (WGS) and whose genomic and annotated clinical data were available through the Hartwig Medical database. Data access was approved by the institute’s Data Access Board (DR-311) and use of data from opt-in routine-care patients by the institutional review board of the Netherlands Cancer Institute. The study followed applicable national requirements and the STROBE reporting guideline.

Adolescents and Young Adults (AYAs) were defined as patients aged 18-39 at diagnosis or biopsy, whereas patients aged 40 or older were classified as older adults (OAs). Age was determined using a tiered approach according to available clinical data. For AYA patients included in an ongoing prospective AYA WGS effort who provided consent for linkage with the Netherlands Cancer Registry (NKR), age at diagnosis was derived from the NKR incidence date. For other patients, age at first systemic treatment or biopsy was used as a proxy to assign age group. Patients who had consented to NKR linkage, but for whom incidence dates could not be successfully linked and age at biopsy was 40 years or older were excluded (n=34). One WGS sample was included per patient. The final study cohort comprised 698 AYAs and 5,220 OAs.

### Cancer type classification and older adult comparison cohorts

Tumors were classified into major cancer groups and more detailed subgroups using the clinical and histopathological annotations available in the Hartwig Medical Database. For primary pan-cancer genomic comparisons, OAs were randomly sampled within each major cancer group to create a 1:1 frequency-matched cohort with the same cancer group-distribution as the AYA cohort. Random sampling was performed using a fixed seed to ensure reproducibility. Subsequent cancer group-specific analyses used all available AYA and OA tumors within each respective cancer group, thereby controlling for differences in overall cancer group composition while maximizing the number of tumors available for the analyses. The complete unmatched cohort of 698 AYAs and 5,220 OAs was additionally analyzed to evaluate the influence of the cohort’s cancer group composition on pan-cancer genomic differences.

### Whole-genome sequencing and genomic annotation

Paired tumor-normal WGS data were processed using the Hartwig Medical Foundation diagnostic bioinformatics pipeline. Somatic single-nucleotide variants (SNVs), multi-nucleotide variants (MNVs), insertions and deletions (indels), copy-number alterations, structural variants, gene fusions, viral integrations, tumor purity, ploidy, whole-genome duplication (WGD), molecular profiles and trinucleotide sequences, and germline variants were obtained from the processed pipeline output. The diagnostic pipeline is open source and available through https://github.com/hartwigmedical/pipeline5.

Somatic driver alterations were obtained from patient-specific driver catalogs generated within the Hartwig pipeline and retained when the pipeline-assigned driver likelihood was ≥0.50. Reported gene fusion events were obtained from the LINX fusion output, and reported viral integrations from the viral annotation output. For prevalence analyses, each gene or rearrangement was counted only once.

Homologous recombination deficiency (HRD) was defined using the CHORD classifier. Microsatellite instability (MSI) was defined from the pipeline’s *MSIsensor*-based classification.

### Somatic genomic landscape analyses

Genome-wide somatic small variant burden was determined by filtering out artifact variants, and quantified separately for SNVs, indels and MNVs. Structural variant junction burden was quantified as the summed number of underlying SV junctions across non-artifact resolved LINX clusters.

Gene-level copy number features were derived from the pipeline’s PURPLE copy number estimates. High-level amplifications were defined as a minimum gene copy number greater than three times the tumor ploidy, and homozygous deletions as a maximum gene copy number below 0.5. Tumor ploidy and WGD status were also obtained from the PURPLE pipeline output.

Pan-cancer somatic driver prevalence was calculated at patient level using mutations, amplifications and deletions. Gene fusions, gene-level disruptive structural variants, and viral integrations were analyzed separately; gene fusion events were obtained from the LINX fusion output and restricted to fusions meeting LINX reportability criteria (reported = TRUE). Gene-level disruptive structural variants were derived from the LINX break-end output using events marked as reported disruptions (reportedDisruption = 1) and counted once per patient-gene pair. For the primary driver landscape, all genes with a significant age-associated prevalence difference after Benjamini-Hochberg correction were displayed and supplemented to a total of 30 genes with the most prevalent remaining drivers across either age group. Statistical testing included the complete set of driver genes.

Protein-level mutation spectra were evaluated for selected recurrent driver genes using reported somatic variants within driver-positive sample-gene pairs.

### Comparison with pediatric cancer drivers

To place the AYA driver landscape in a broader age context, gene-level driver prevalence was compared with pediatric cancer using data from the PeCan resource of the St. Jude Children’s Research Hospital (https://pecan.stjude.cloud/variants/prevalence/, accessed 21 September 2026). Analyses were restricted to soft tissue tumors and CNS tumors, for which sufficiently comparable pediatric and adult cancer samples were available. Pediatric prevalence was defined as the proportion of cases with at least one reported variant in the respective gene, as provided by PeCan. Because the PeCan and Hartwig datasets use different analytical frameworks, these comparisons were considered descriptive.

For the Hartwig AYA and OA cohorts, gene-level prevalence was calculated from somatic driver mutations, amplifications, deletions, and gene fusions using all available tumors within the corresponding cancer group. Genes displayed in the visualization represent the union of the 20 most prevalent genes in each age group. Genes were grouped according to their age-associated prevalence pattern, as pediatric-predominant, shared pediatric/AYA, AYA-predominant, AYA-depleted, shared AYA/OA, OA-predominant, or shared across ages. Adjacent age groups were considered to show a shared predominant pattern when their prevalences differed less than 1.5-fold. Pattern assignment used unrounded prevalence estimates for AYAs and OAs, and the whole-number prevalences as reported by PeCan for pediatric tumors.

### Assignment and analysis of mutational signatures

Matrices containing the count of mutations at classified context categories as defined in the COSMIC catalog were generated using the R package *mutSigExtractor* (v1.29), available via https://github.com/UMCUGenetics/mutSigExtractor. Separate mutation matrices for single-base substitution (SBS), insertion or deletion (ID), and doublet-base substitution (DBS) categories were generated for each major primary tissue category, focusing on the tissue groups with more than 20 cases for both OA and AYA patients.

For each tissue-specific mutation matrix, mutational signatures of the COSMIC catalog v3.5 were assigned to the SBS, ID and DBS mutation profiles of all AYA and OA samples using *SigProfilerAssignment* (v1.1.3),^12^ which mathematically resolves the lowest number of signatures providing the best fit to the mutational profile of a sample, resulting for each sample a set of signatures and their respective contribution. Due to this signature selection, most signatures in most samples will have zero contribution. For refitting across all three mutation types, *SigProfilerAssignment* was run with the setting to remove all signatures classified as artifact: ‘exclude_signature_subgroups = Artifact_signatures’.

To determine whether specific samples, or specific subgroups of samples were not adequately refitted we employed a cosine similarity between the reconstructed mutational profile and the original mutation profile across the samples (displayed in Extended Data Fig. 5) using the ‘cos_sim’ function from the R package *MutationalPatterns* (v3.22.0). High cosine similarity indicates good matching of the contributing signatures from the COSMIC (v3.5) catalog to the existing patterns.

Using mutational signature activity as a proxy for mutational processes, we sought to find significant differences in signature contributions across AYA and OA samples. To compare contribution, which follows a non-normal distribution, we employed a Wilcoxon rank-sum test on the absolute contribution of mutations of each sample, correcting for p-value using FDR. Significant enrichments between signature presence or absence across AYA and OA cohorts for specific signatures were determined using a Fisher’s exact test.

### Pathogenic germline variant assessment

Reported germline findings in AYAs and the 1:1 cancer group-matched OA cohort were expert-reviewed by a certified clinical laboratory geneticist. The investigated genes were based on the list of genes considered for clinical genetics referral in the Netherlands, with additional cancer predisposition genes considered relevant to the study. Findings were retained when classified as pathogenic or likely pathogenic, or when manual review supported inclusion based on the available molecular and clinical context. For each retained finding, tumor type-concordance was assigned according to whether the affected gene was associated with the patient’s tumor type or subtype, based on the guideline of the Dutch Society for Clinical Genetics, the Dutch Society for Pathology, and the Dutch Association of Clinical Genetic Laboratory Diagnostics.^10,16^ Findings outside the expected tumor context were retained and analyzed separately as non-concordant findings. Analyses were performed at the patient-gene level, with each affected gene counted once per patient.

For recessive cancer predisposition genes, including *MUTYH*, *NTHL1*, *RAD50* and *FANCL*, monoallelic findings were not considered relevant unless homozygous or compound heterozygous germline alterations were present, or the combined germline and tumor context supported functional biallelic inactivation.

### Genomic second hit assessment and tumor genomic context

For each retained germline finding, tumor involvement of the same gene was assessed using somatic SNVs or indels, loss of heterozygosity (LOH) or tumor-biallelic loss, and structural variants affecting the same gene. Because allelic phasing between germline and somatic alterations was generally unavailable, these events were considered putative genomic second hits. A genomic second hit was assigned when at least one qualifying same-gene somatic event, tumor-biallelic event, LOH event, or disruptive structural variant was identified. HRD and MSI presence were evaluated separately and not visualized as genomic second hit in the germline figures.

Gene-level LOH was defined as a minor allele copy number <0.5. LOH was considered supportive of a genomic second hit only when the tumor copy number context supported retention of the variant allele, which was defined as a variant copy number >0.3. LOH with variant copy number ≤0.1 was considered compatible with loss of the germline variant allele, and therefore not counted as a second hit. Ambiguous LOH events were not included as genomic second hits.

To assess the functional genomic context of inherited DNA-repair defects, PGVs in homologous recombination genes (*BRCA1*, *BRCA2*, *PALB2*, and *RAD51C*) were integrated with the pipeline’s CHORD-defined HRD status and with SBS3/ID6 contributions from the signature refitting analyses. PGVs in mismatch repair genes (*MLH1*, *MSH2*, *MSH6*, *PMS2*, and *EPCAM*) were integrated with the pipeline’s MSIsensor-based MSI status and with the summed MMR-associated SBS (SBS6/SBS14/SBS15/SBS20/SBS21/SBS26/SBS44) and ID (ID1/ID2/ID7) burdens from the signature refitting analyses. Biallelic germline alterations, including compound heterozygous variants, were considered separately from acquired genomic second hits.

### Statistical analysis

Continuous variables were summarized using medians and interquartile ranges unless otherwise specified and compared between age groups using two-sided Wilcoxon rank-sum tests. Categorical variables and differences in feature prevalence were assessed using Fisher’s exact tests. For analyses involving multiple genomic features, p-values were adjusted using Benjamini-Hochberg false discovery rate (FDR) correction within each respective analysis. Nominal p-values were used for prespecified single-feature comparisons unless otherwise indicated. Statistical significance was defined as p<0.05 or FDR-adjusted p<0.05, as applicable.

Analyses were performed at patient level using one WGS sample per patient. Cancer group-specific analyses included all available tumors within the respective cancer group, whereas primary pan-cancer comparisons used the cancer group-matched cohort. Missing data were not imputed. Analyses were performed in R version 4.4.1 using Jupyter notebooks.

## ARTIFICIAL INTELLIGENCE STATEMENT

During preparation of this manuscript, ChatGPT (GPT-5.6 Sol; OpenAI; September 2026) was used for copy-editing and troubleshooting during code development. All artificial intelligence-assisted outputs were critically reviewed and revised by the authors, who take full responsibility for the accuracy and integrity of the manuscript and analyses.

## DATA AVAILABILITY

Raw and processed WGS data from the Hartwig Medical database are available for academic research under controlled access through Hartwig Medical Foundation’s data access request procedures, subject to Data Access Board review and a data transfer agreement. The database identifiers for the study cohort are provided in Supplementary Table 1 and can be used to request the corresponding genomic and linked clinical data.

Requests can be submitted to. Detailed procedures, guidelines and application forms are available through https://www.hartwigmedicalfoundation.nl/en/data/data-access-request/.

## CODE AVAILABILITY

Custom code used for data preparation, statistical analyses, mutational signature extraction and refitting, and figure generation will be made available upon publication; The diagnostic WGS bioinformatics pipeline used by Hartwig Medical Foundation is open source and available at https://github.com/hartwigmedical/pipeline5/.

## LITERATURE

1. Berkman, A. M., Mittal, N. & Roth, M. E. Adolescent and young adult cancers: unmet needs and closing the gaps. Current Opinion in Pediatrics 35, 84–90 (2023).

2. Hughes, T., et al. The current and future global burden of cancer among adolescents and young adults: a population-based study. The Lancet Oncology 25, 1614–1624 (2024).

3. on behalf of the Biology and Clinical Trials Subgroups of the US National Cancer Institute Progress Review Group in Adolescent and Young Adult Oncology et al. The distinctive biology of cancer in adolescents and young adults. Nat Rev Cancer 8, 288–298 (2008).

4. Van Putten, J., et al. Genomic distinctions in adolescent and young adult cancer: A comprehensive review. Cancer Treatment Reviews 140, 103021 (2025).

5. Hayashi, N., et al. Addressing the knowledge gap in the genomic landscape and tailored therapeutic approaches to adolescent and young adult cancers. ESMO Open 9, 103659 (2024).

6. Wang, X., Langevin, A.-M., Houghton, P. J. & Zheng, S. Genomic disparities between cancers in adolescent and young adults and in older adults. Nat Commun 13, 7223 (2022).

7. Mosele, M. F., et al. Recommendations for the use of next-generation sequencing (NGS) for patients with advanced cancer in 2024: a report from the ESMO Precision Medicine Working Group. Annals of Oncology 35, 588–606 (2024).

8. Mandelker, D., et al. Germline-focussed analysis of tumour-only sequencing: recommendations from the ESMO Precision Medicine Working Group. Annals of Oncology 30, 1221–1231 (2019).

9. Roepman, P., et al. Clinical Validation of Whole Genome Sequencing for Cancer Diagnostics. The Journal of Molecular Diagnostics 23, 816–833 (2021).

10. Koster, R., et al. Impact of genetic counseling strategy on diagnostic yield and workload for genome-sequencing-based tumor diagnostics. Genetics in Medicine 26, 101032 (2024).

11. Martínez-Jiménez, F., et al. Pan-cancer whole-genome comparison of primary and metastatic solid tumours. Nature 618, 333–341 (2023).

12. Díaz-Gay, M., et al. Assigning mutational signatures to individual samples and individual somatic mutations with SigProfilerAssignment. Bioinformatics 39, btad756 (2023).

13. Rosendahl Huber, A., et al. Improved detection of colibactin-induced mutations by genotoxic E. coli in organoids and colorectal cancer. Cancer Cell 42, 487–496.e6 (2024).

14. Díaz-Gay, M., et al. Geographic and age variations in mutational processes in colorectal cancer. Nature 643, 230–240 (2025).

15. Chowdhury, T. et al. Tumor-initiating genetics and therapy drive divergent molecular evolution in IDH-mutant gliomas. bioRxiv 10.1101/2025.07.11.664189 (2025) doi:10.1101/2025.07.11.664189.

16. Projectgroep tumor-en erfelijkheidsdiagnostiek van de NVVP, VKGL en VKGN. [Table 3 Guideline for reporting of advices concerning germline diagnostics in pathology reports and during MTB discussions about the analysis of solid tumours in adult cancer patients]. (2023).

17. Luen, S. J., et al. Genomic characterisation of hormone receptor-positive breast cancer arising in very young women. Annals of Oncology 34, 397–409 (2023).

18. Serio, P. A. D. M. P., et al. Somatic Mutational Profile of High-Grade Serous Ovarian Carcinoma and Triple-Negative Breast Carcinoma in Young and Elderly Patients: Similarities and Divergences. Cells 10, 3586 (2021).

19. Lim-Fat, M. J., et al. A comparative analysis of IDH-mutant glioma in pediatric, young adult, and older adult patients. Neuro-Oncology 26, 2364–2376 (2024).

20. Wu, J., Heidelberg, R. E. & Gajjar, A. Adolescents and Young Adults With Cancer: CNS Tumors. JCO 42, 686–695 (2024).

21. Van Der Graaf, W. T. A., Orbach, D., Judson, I. R. & Ferrari, A. Soft tissue sarcomas in adolescents and young adults: a comparison with their paediatric and adult counterparts. The Lancet Oncology 18, e166–e175 (2017).

22. Debortoli, E., et al. Utility of genomic testing in children, adolescents, and young adults with cancer. JNCI: Journal of the National Cancer Institute djae233 (2024) doi:10.1093/jnci/djae233.

23. Cercek, A., et al. A Comprehensive Comparison of Early-Onset and Average-Onset Colorectal Cancers. JNCI: Journal of the National Cancer Institute 113, 1683–1692 (2021).

24. Toss, A., et al. Cancer Predisposition Genes in Adolescents and Young Adults (AYAs): a Review Paper from the Italian AYA Working Group. Curr Oncol Rep 24, 843–860 (2022).

25. Waks, A. G., et al. Somatic and Germline Genomic Alterations in Very Young Women with Breast Cancer. Clinical Cancer Research 28, 2339–2348 (2022).

26. Priestley, P., et al. Pan-cancer whole-genome analyses of metastatic solid tumours. Nature 575, 210–216 (2019).

27. Langenberg, K. P. S., et al. Implementation of paediatric precision oncology into clinical practice: The Individualized Therapies for Children with cancer program ‘iTHER’. European Journal of Cancer 175, 311–325 (2022).

28. Hodder, A., et al. Benefits for children with suspected cancer from routine whole-genome sequencing. Nat Med 30, 1905–1912 (2024).

29. Van Putten, J., et al. Real-world clinical utility of tumor whole-genome sequencing in solid cancers. Nat Med 32, 1286–1295 (2026).

